# Effects of Peripheral Haptic Feedback on Intracortical Brain-Computer Interface Control and Associated Sensory Responses in Motor Cortex

**DOI:** 10.1101/2020.08.17.240648

**Authors:** Darrel R. Deo, Paymon Rezaii, Leigh R. Hochberg, Allison M. Okamura, Krishna V. Shenoy, Jaimie M. Henderson

## Abstract

Intracortical brain-computer interfaces (iBCIs) provide people with paralysis a means to control devices with signals decoded from brain activity. Despite recent impressive advances, these devices still cannot approach able-bodied levels of control. To achieve naturalistic control and improved performance of neural prostheses, iBCIs will likely need to include proprioceptive feedback. With the goal of providing proprioceptive feedback via mechanical haptic stimulation, we aim to understand how haptic stimulation affects motor cortical neurons and ultimately, iBCI control. We provided skin shear haptic stimulation as a substitute for proprioception to the back of the neck of a person with tetraplegia. The neck location was determined via assessment of touch sensitivity using a monofilament test kit. The participant was able to correctly report skin shear at the back of the neck in 8 unique directions with 65% accuracy. We found motor cortical units that exhibited sensory responses to shear stimuli, some of which were strongly tuned to the stimuli and well modeled by cosine-shaped functions. We also demonstrated online iBCI cursor control with continuous skin-shear feedback driven by decoded command signals. Cursor control performance increased slightly but significantly when the participant was given haptic feedback, compared to the purely visual feedback condition.

## 1 INTRODUCTION AND BACKGROUND

**P**EOPLE coordinate, plan, and execute movements under the guidance of both proprioception and vision [1]–[3], which are vital in informing an estimate of the body’s configuration and motion in space. When there is a deficit or loss in proprioception, simple motor control and dexterous object manipulation becomes disrupted and uncoordinated [4]–[9]. In the case of most intracortical brain-computer interface (iBCI) users, this proprioceptive deafferentation is additionally accompanied by the lack of physical motion when attempting movement. Thus, iBCI users rely heavily on visual feedback alone when performing control tasks. In order to achieve naturalistic movement and function similar to native limbs, iBCI systems will likely need to include somatosensory feedback as a means to provide artificial proprioception.

Recently, there has been an increased effort to develop bidirectional neural interfaces capable of both measuring neural signals from the brain and providing sensory signals back to the user [10]–[12]. Predominantly, intracortical microstimulation (ICMS) – electrically stimulating the cortex – has been used to artificially evoke both tactile and proprioceptive percepts in nonhuman primates (NHPs) and people. Studies have shown that ICMS of the primary sensory cortex (S1) has enabled NHPs to perform sensory discrimination tasks with performance similar to mechanical stimulation of the hand [13]–[16]. Additionally, ICMS has been shown to effectively communicate task-relevant feedback signals that guide online, multidimensional movement control in NHPs, acting as a form of artificial proprioception that NHPs are capable of learning [17]. More recent ICMS studies in people have elucidated characteristics of stimulation-evoked sensations and report both tactile [10] and proprioceptive [12] percepts. Although ICMS of S1 cannot perfectly mimic natural sensory percepts, people can learn to use the evoked percepts as feedback for improved neuroprosthetic control [18].

With the development of bidirectional neural interfaces, it is also important to consider the effects of sensory stimulation on the motor cortex. The primary motor cortex (M1) has been shown to be responsive to many types of sensory inputs, including visual, tactile, and proprioceptive [19]. Previous single-unit electrophysiology studies in NHPs showed that some M1 cortical units are responsive to tactile stimulation, as well as active and passive movement of the limbs [20], [21]. More recently, a study measuring neural activity from electrocorticography (ECoG) grids placed on the “hand” area of the human motor cortex of people undergoing invasive monitoring for epilepsy indicated neural responses to passive tactile stimulation of the palm [22]. However, when considering iBCI users with impaired sensory pathways (e.g., spinal cord injury), tactile stimulation may need to be provided on areas of the body that do not necessarily correspond with the areas of the brain used for control.

More recently, we have found that the hand knob area of premotor cortex (dorsal precentral gyrus) in people with tetraplegia, including the participant mentioned in this study, is involved with movements spanning the entire body and not just limited to movements involving the arm and hand [23]. Specifically, we found that overt or attempted movements of the face, head, leg, and arm modulated neural activity. Considering the intermixed whole-body tuning of this small patch of premotor cortex – contrary to traditional expectations of macroscopic somatotopy as proposed by the motor homunculus [24] – it is prudent to also consider that the sensory homunculus analog may have similar whole-body representation where tactile stimulation on the neck may result in neural activity in areas of the sensory cortex outside of the conventional neck/head area. This may in turn lead to responses in M1 cortical units at the site of recording.

Here, we integrated a commercially available haptic device into our existing iBCI system, which provided skin-shear haptic stimulation at the back of the neck in a research participant with an iBCI system. Stimulation was delivered at a location which specified by assessing sensitivity to touch using a clinical monofilament test kit. Next, we assessed perception of shear in 8 radial directions. Our participant was able to verbally discriminate the 8 shear directions with an accuracy of 65.6%. In addition, we found motor cortical units that exhibited sensory responses to the shear stimuli, some of which were strongly tuned to the stimuli and well modeled by cosine-shaped functions. Finally, we demonstrated online iBCI cursor control with continuous skin-shear feedback driven by decoded command signals, and compared performance to a purely visual feedback condition.

## 2 METHODS

### 2.1 Study Permissions and Participant Details

A single participant (T5) enrolled in the BrainGate2 Neural Interface System clinical trial (ClinicalTrials.gov Identifier: NCT00912041, registered June 3, 2009) gave informed consent prior to this study. This pilot clinical trial was approved under an Investigational Device Exemption (IDE) by the US Food and Drug Administration (Investigational Device Exemption #G090003). Permission was also granted by the Stanford University Institutional Review Board (protocol #20804).

Participant T5 is a right-handed man (65 years of age at the time of study) with tetraplegia due to cervical spinal cord injury (classified as C4 AIS-C) that occurred approximately 9 years prior to study enrollment. T5 was implanted with two 96-electrode (1.5 mm length) intracortical micro-electrode arrays (Blackrock Microsystems, Salt Lake City, UT) in the hand knob area of the left (dominant) precentral gyrus. T5 retained full movement of the face and head and the ability to shrug his shoulders. Below the level of his spinal cord injury, T5 retained some very limited voluntary motion of the arms and legs that was largely restricted to the left elbow. We refer to any intentional movements of the body below the level of injury as being ‘attempted’ movements.

### 2.2 Neural Signal Processing

Neural signals were recorded from the microelectrode arrays using the NeuroPortTM system from Blackrock Microsystems (Hochberg et al. [25] describes the basic setup). Neural signals were analog filtered from 0.3 Hz to 7.5 kHz and subsequently digitized at 30 kHz (with 250 nV resolution). The digitized signals were then sent to control computers for real-time processing to implement the iBCI. The real-time iBCI was implemented in custom Simulink Real-Time software running on a dedicated PC with the xPC real-time operating system (Mathworks Inc., Natick, MA).

To extract action potentials (spikes), the signal was first common-average re-referenced within each array. Next, a digital bandpass filter from 250 Hz to 3 kHz was applied to each electrode before spike detection. For threshold crossing detection, we used a −4.5 x RMS threshold applied to each electrode, where RMS is the electrode-specific root-mean-square of the voltage time series recorded on that electrode. In keeping with standard iBCI practice, we did not spike sort (i.e., assign threshold crossings to specific single neurons) [26]–[29].

### 2.3 Cutaneous Sensitivity Testing

We used a Semmes-Weinstein monofilament examination (SWME) kit (North Coast Medical, Inc., Gilroy, CA) to evaluate cutaneous sensation levels on the body to identify locations to provide haptic feedback during iBCI control. The SWME kit includes a set of handheld monofilament probes, each calibrated within a 5% standard deviation (SD) of their target force level. A monofilament is pressed into the skin at a test site perpendicular to the skin surface until it bends when the peak force reaches the target threshold, maintaining the target force under bending.

In addition to T5, we recruited an able-bodied control group of adult participants comprising 3 males 28±3 (mean ± SD) years of age and 7 females 27±6 years of age. Participants in the control group had no known sensory impairment or loss. The experimental protocol for the cutaneous sensitivity test in healthy participants was approved by the Stanford University Institutional Review Board (protocol #22514) and all participants gave informed, written consent prior to participation.

All participants were required to keep their eyes closed throughout the exam. Probing locations were selected at random from a predetermined set of locations (Fig. 4). The total set of probing locations was determined after pilot sensitivity tests with T5 indicated that his sensitivity to touch below the neckline, including areas of the arm and hand, was greatly degraded due to the level of spinal cord injury. T5 was unable to feel any of the monofilament probes at the hands or arms. Participants were not provided with any knowledge of the probing locations prior to the exam and were only informed that areas of the upper body could potentially be probed. A probe consisted of slowly pressing the filament (always starting with the filament size marked with unitless label ‘2.83’, which is categorized as healthy touch sensitivity) against the skin until bending. The probe was held in the bent state for approximately 1.5 seconds and then removed. This was repeated for the same filament until either a verbal response was elicited or a total of three probes occurred. If a response was elicited, we proceeded to the next smallest filament size, repeating the process until the participant was unable to feel the probe, at which point the last filament size and associated probing force to elicit a response was recorded for that location. If a response was not elicited with the initial filament size of 2.83, the next largest filament size was selected and the probing sequence repeated.

For the control group, average sensitivity at each probing location was computed via the sample mean and 95% confidence intervals were computed using bootstrap resampling (100,000 iterations). For T5, there was only a single data point for each probed location.

### 2.4 Perception of Skin-Shear on Back of Neck

#### 2.4.1 Haptic Device Hardware and Software

We used a Phantom Premium 1.5 (3D Systems, Inc., Rock Hill, SC) haptic device, which has been primarily used in haptic and telerobotic applications [30]–[32]. We positioned a Premium on top of a table behind participant T5, with the linkages oriented such that the stylus end-effector’s axial axis was perpendicular to the plane of the back of the neck, as shown in Figure 1. A Nano17 (ATI Industrial Automation, Inc., Apex, NC) 6-axis force sensor was attached to the end of the stylus with a custom 3D-printed adapter. We fixed a tactor to the free end of the force sensor, which was covered with a piece of double-sided tape designed for adhesion to the skin (3MTM, Santa Clara, CA). At the start of each session, we marked the desired tactor contact position on the back of T5’s neck with a marker and cleaned the area with an alcohol pad. Next, a fresh piece of tape was applied to the tactor surface. Finally, the tactor was carefully positioned and pressed into the skin at the desired location for approximately 10 seconds. Adhesion was tested by vigorously moving the stylus by hand in all directions and ensuring no slip of the tactor along the skin. T5’s head rested on the wheelchair’s headrest to ensure minimal movement of the neck and head against the tactor.

**Fig. 1:**
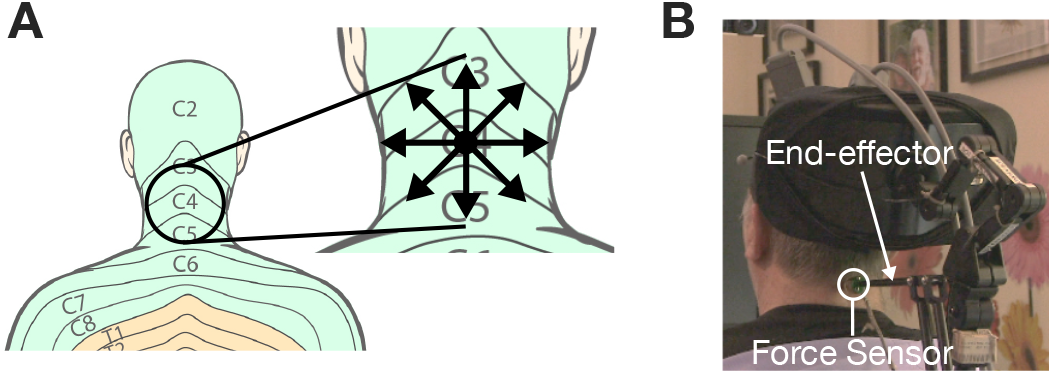
Skin-shear haptic stimulation on back of neck. (A) Target stimulus location for participant T5 is at the center of the C4 dermatome. The inlay illustrates the 8 radial directions along which shear force was provided during the perception study. (B) Actual photograph of the haptic device configuration. Dermatome images adapted from Janet Fong [33].

The haptic device was controlled by a separate computer that also logged force sensor measurements by custom C/C^++^ software developed in the Microsoft Visual Studio IDE. The CHAI3D open source framework [34] was used to render haptic interaction at a control-loop rate between 4 and 9 kHz (which is the native CHAI3D haptic thread rate range). All other loop rates (e.g., main state machine, data logging, etc.) operated at 1 kHz. Synchronization and communication between the haptic device control computer and iBCI control computer was facilitated by UDP – low-latency and loss-tolerating communication – and timestamping through a wired ethernet connection to a local area network hub.

#### 2.4.2 Perception Task

Prior to assessing T5’s perception of shear force on the back of the neck, we conducted a pilot study to determine the range of shear force parameters (magnitude and direction) to probe. We found that a normal force of approximately 0.3 N into the skin surface was sufficient to maintain contact between the tactor and the neck site for shear forces up to a maximum of 1 N. To prevent slip of the tactor from the neck site, we limited the maximum shear force in any direction to 0.5 N, which was detectable by T5.

The focus on shear direction was motivated by previous iBCI cursor control studies suggesting that when a neurally-controlled cursor is far from the target, the normalized neural population activity is similar to a unit vector pointing straight from the cursor to the target [35]–[37]. A more recent study found that iBCI users apply a diminishing ‘neural push’ to the cursor as it approaches the target [38]. Considering the initial, or ballistic, movement out to a target, we can approximate the ‘neural push’ as being saturated in magnitude and pointing in the direction of the target. Hence, direction information may be most useful during the initial parts of movement.

We performed a perception study across two consecutive session days. For each session, T5 was asked to close his eyes and focus on the haptic stimulation provided on the back of his neck. The haptic device was configured as pictured in Figure 1B where the tactor was pressed into the neck with a normal force of 0.3 N. T5 was instructed to report the direction in which he felt the haptic stimulus. His preferred reporting method was referencing the face of a clock. T5 was not given any information regarding the stimuli directions and was unaware that 8 unique directions were being tested. T5 was asked to be precise in his reporting. A block consisted of approximately 244 trials (a total of 488 trials over the course of two sessions). A shear stimulus in one of eight directions (Fig. 1A) was presented pseudo-randomly (i.e., randomly within sets of 8 where each set contained exactly one of each stimulus) to ensure a balanced number of repetitions per each unique stimulus. Each trial was 4 seconds in total duration. After an idle period of 1 second, a nominally 0.5 N shear force stimulus was provided as a step input for a length of 1 second, followed by another idle period of 2 seconds. Thus, there were exactly 3 seconds between each stimulus.

### 2.5 Tactile Sensory Responses in Motor Cortex

#### 2.5.1 Study Structure

The design of this study was similar to the perception study discussed prior, except T5 was asked to not verbally respond and remain in an idle state. An idle state was defined as not attempting any movements, not imagining any movements, and not thinking about anything in particular. He was urged to not attend to the stimulus in any manner throughout the study. T5 was not given any information about the stimuli or timing of the study. We conducted two five-minute blocks (60 trials each) where a shear stimulus in one of eight radial directions (Fig. 1A) was presented pseudo-randomly (random within sets of 8, where each set contained exactly one of each stimulus).

#### 2.5.2 Neural Data Analysis

Spike times were separated into 10 ms bins and z-scored. Z-scoring was accomplished by first subtracting, in a block-wise manner, the mean spike count over all 10 ms bins within each block. After mean subtraction, the binned spike counts were divided by the sample standard deviation computed using all 10 ms bins across all blocks. Electrodes with firing rates less than 1 Hz over all time steps were excluded from further analysis to de-noise population-level results. To visualize an electrode channel’s firing rate responses to the sensory stimulation, spike trains were smoothed with a Gaussian kernel (with 30 ms standard deviation) to reduce high frequency noise.

To assess neural tuning to sensory shear stimulation on a given electrode, we used a 1-way ANOVA (*α* = 0.05) of firing rates observed during sensory stimulation and firing rates observed during idle. We first computed firing rates for each trial within the 500 to 900 ms window relative to stimulus onset, and within the −400 to 0 ms window relative to stimulus onset to represent ‘idle’ or ‘baseline’ activity. Next, we grouped each of the computed firing rates into either their respective stimulus directions or the additional ‘baseline’ group and performed a 1-way ANOVA. If the p-value was less than 0.001, the electrode was considered to be strongly tuned to skin-shear stimulation on the back of the neck.

To assess the tuning strength of each strongly tuned electrode to shear stimulation, we computed FVAF (fraction of variance accounted for) scores [23]. The FVAF score was computed using the following equations:

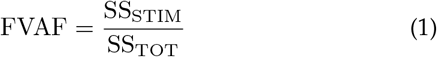

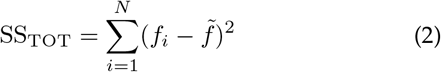

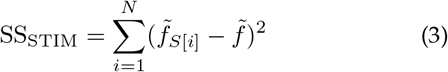

Here, SS_TOT_ is the total variance (sum of squares), SS_STIM_ is the shear stimulation-related variance, *N* is the total number of trials, *f_i_* is the firing rate for trial *i* within the 500 to 900 ms window after stimulus onset, 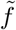 is the average firing rate within the window across all shear directions, and 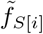 is the average firing rate for the particular stimulus cued on trial *i*. FVAF scores range from 0 (no stimulation-related variance) to 1 (all variance is stimulation-related).

To characterize the stimulation-related responses in the measured motor cortical units, we fit tuning curves to each of the strongly tuned electrode channels. A tuning curve relates the firing rate of a particular channel to a presented stimulus. Tuning curves were fit to the following function:

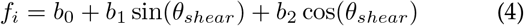

Here, parameter *b*_0_ represents the baseline firing rate, parameter *b*_1_ is the *y* component of the preferred stimulus direction, parameter *b*_2_ is the *x* component of the preferred stimulus direction, *f_i_* is electrode *i*’s firing rate within the 500 to 900 ms window, and *θ_shear_* is the angle of the shear stimulus in radians. Goodness-of-fit-adjusted R-squared statistics were computed as indicators of the fit quality of each tuning curve. The adjusted R-squared statistic can take on values less than or equal to 1, with a value closer to 1 indicating a better fit. Negative values of the adjusted R-squared statistic are also possible, indicating that a linear fit is better than the cosine-like tuning model.

To further analyze the stimulus response-related information content encoded in the neural activity, a Naïve Bayes decoder with leave-one-out cross-validation was used to classify the shear stimulus direction on a given trial. The inputs to the decoder were the firing rates computed within the 500 to 900 ms window after stimulus onset. Only strongly tuned channels were used for decoding.

### 2.6 Haptic Feedback During iBCI Training and Control

#### 2.6.1 Integrated iBCI Haptic System

The Phantom Premium haptic device was integrated into the existing iBCI system to facilitate artificial proprioceptive feedback during cursor control (Fig. 2). Artificial proprioceptive feedback has been previously demonstrated in ICMS studies by driving electrical stimulation patterns as functions of iBCI decoded parameters, i.e., mapping decoded velocity signals to stimulation signals [17]. In an effort to mimic this feedback paradigm, we decided to drive the haptic device with our iBCI’s decoded velocity commands, mapping a 2-dimensional decoded velocity vector to a shear force vector on the back of the neck.

**Fig. 2:**
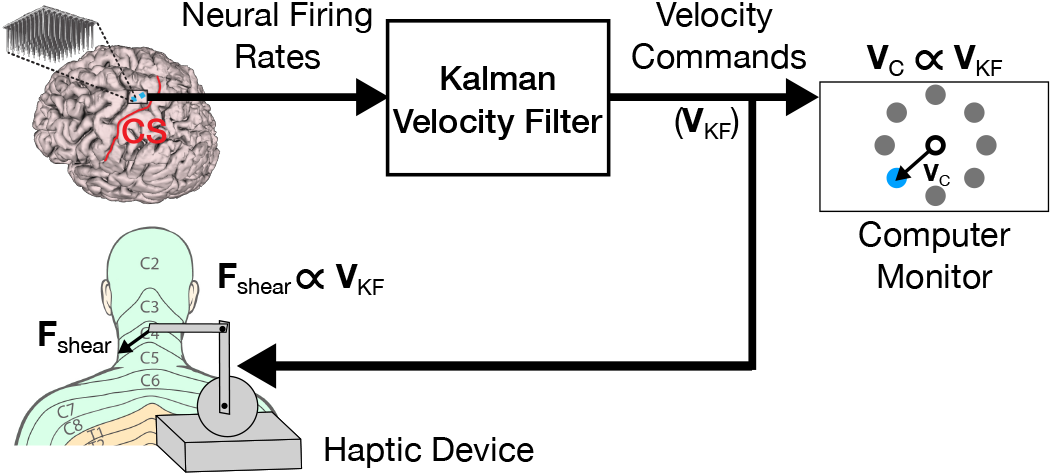
Artificial proprioception via skin-shear haptic feed-back. Neural firing rates are measured and translated to a two-dimensional velocity command vector **V**_KF_ by a Kalman filter. The **V**_KF_ command vector simultaneously drives the velocity of a virtual cursor (**V**_C_) on a computer monitor, and the shear force (**F**_shear_) produced by a haptic device on the back of the participant’s neck. Dermatome image adapted from Janet Fong [33].

In defining the velocity-shear mapping function, we had already established a range of shear force magnitudes to apply (0 to 0.5 N) from pilot and perceptual tasks described earlier. To get a sense of the range and frequency of velocity command values, we visualized the distribution of velocity commands during a typical iBCI cursor control task with no haptic feedback. Figure 3A depicts the distribution of velocity commands in both the X and Y directions for a 10-minute closed-loop block (without haptic feedback condition) from the cursor control task used to assess performance. The units of velocity commands are reported in ‘workspace width per second’ (WW/s). Instead of capturing the entire range of velocity command values, we decided to capture approximately 95% of values about the mean (between −0.35 and 0.35 WW/s) and linearly map that range to the predetermined shear force range. This was done to simplify the mapping function and to capture a symmetric range of most velocity values centered about the mean. This implies that velocity command magnitudes greater than 0.35 WW/s are mapped to the saturated maximum shear force magnitude of 0.5 N. The velocity-shear mapping function was governed by the following piecewise function:

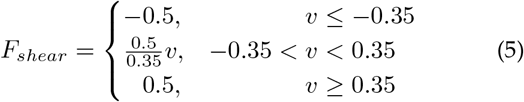

**Fig. 3:**
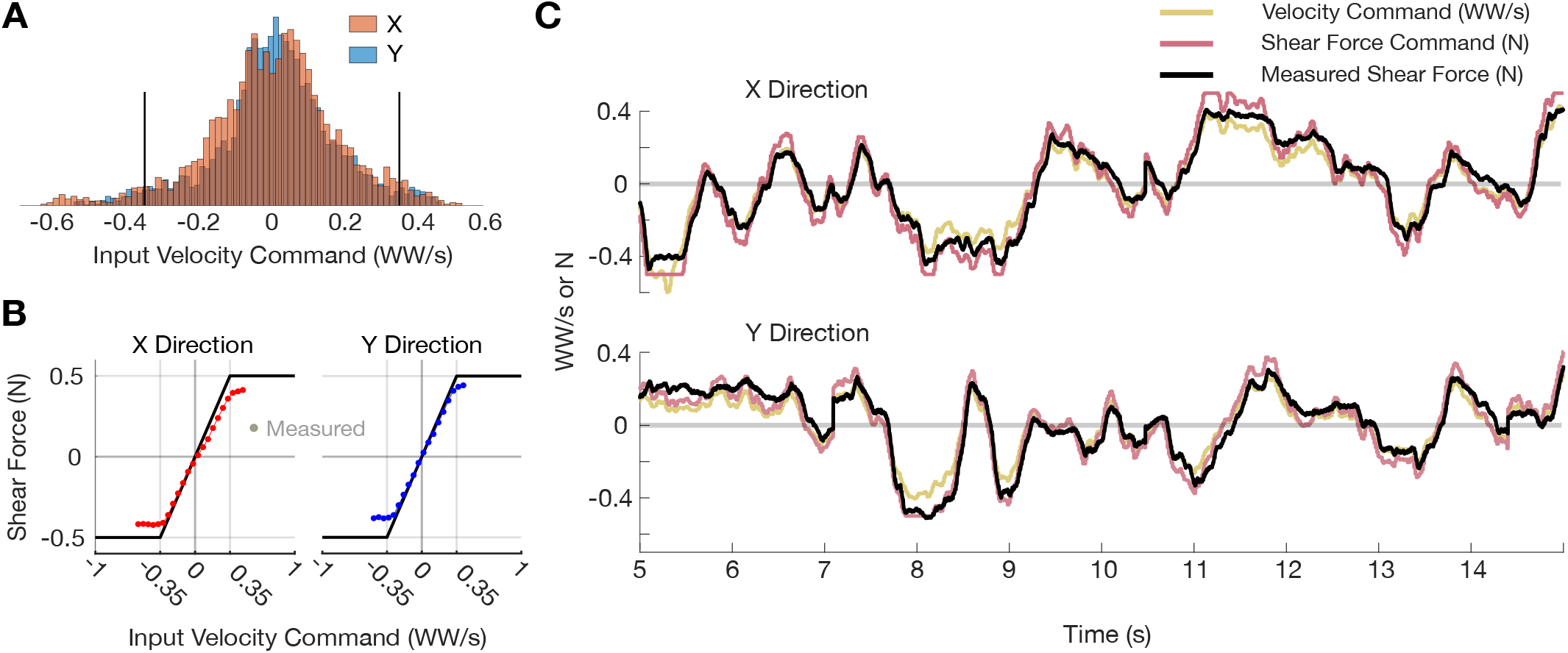
Velocity-shear mapping function and force tracking. (A) Distribution of X- and Y-direction velocity commands from a typical closed-loop iBCI cursor control task. Units are in workspace width per second (WW/s). Black lines bound the 95th percentile (0.35 WW/s). (B) Velocity-shear saturation function (black line) with average measured force plotted as a function of velocity command for the X (red) and Y (blue) directions. The 95% CIs are plotted but are smaller than the width of the plotted points. Data is from a 10-minute iBCI cursor control block with haptic feedback. (C) Force tracking example from a 10 second snippet of data from the task used in panel B. Velocity commands (gold) are mapped to shear force commands (purple) using the function in panel B. The black trace is measured force sensor data.

**Fig. 4:**
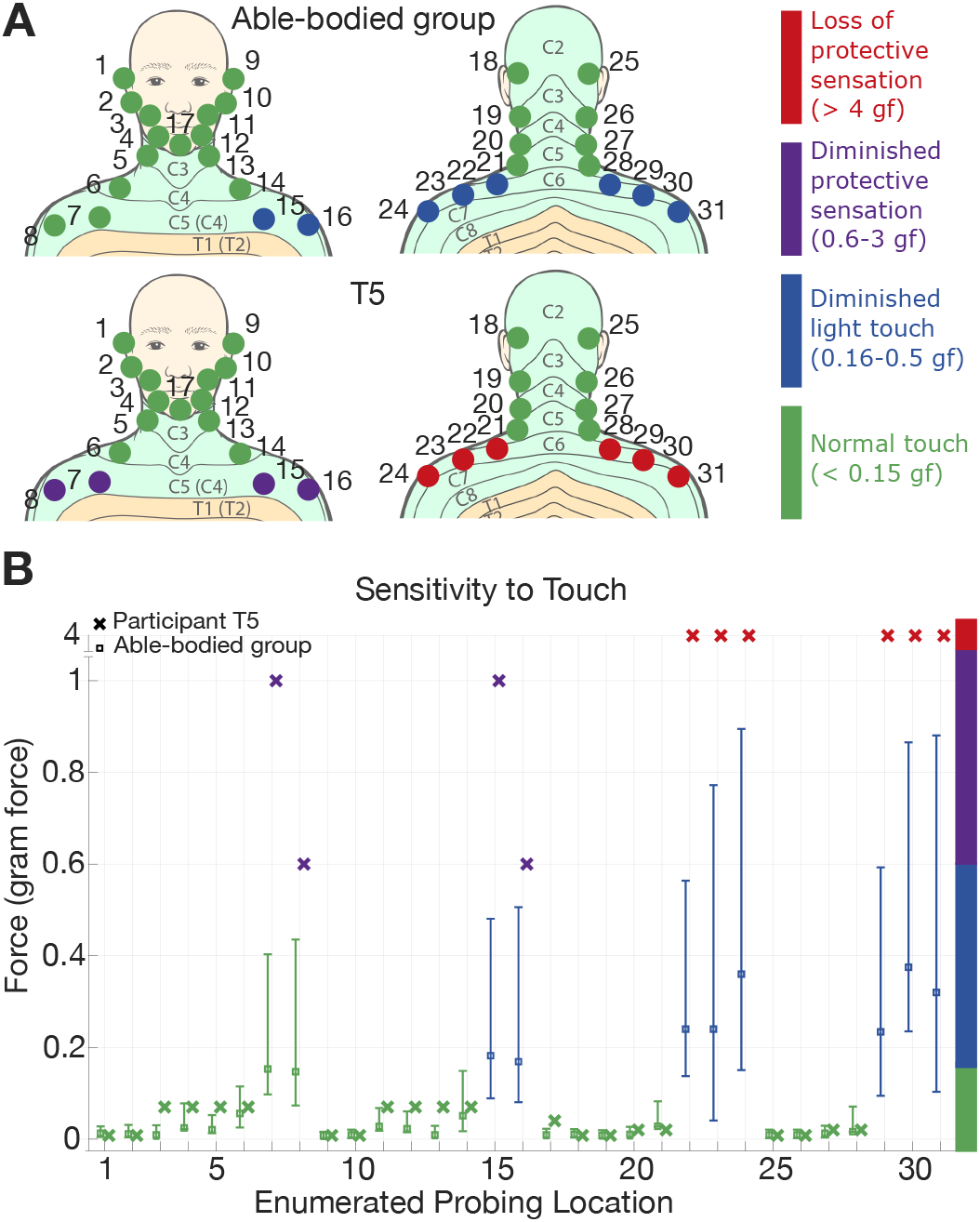
Sensitivity to touch. (A) Able-bodied group average and T5’s sensitivity with corresponding classification colors. (B) Means and 95% CIs for sensitivity (gram-force) at each probing location for the able-bodied group (D) and T5’s responses (**x**). Dermatome images adapted from Janet Fong [33].

Here, *F_shear_* is the desired shear force (units of Newtons) at the end-effector of the haptic device and *v* is the input velocity command (units of WW/s) received over UDP from the iBCI control computer.

The velocity-shear saturation function is depicted in Fig. 3B with averaged measured force sensor data overlaid. Using data from a 10-minute iBCI cursor control block with skin-shear haptic feedback governed by the aforementioned velocity-shear mapping function, we binned the range of velocity commands in 0.05 WW/s bins and computed the mean and 95% confidence intervals (CIs) for the force sensor values measured within each respective bin by fitting to a normal distribution. The average measured force tracks the desired saturation function well, although it begins to saturate at approximately a force magnitude of 0.45 N instead of the desired 0.5 N. This discrepancy may be due to either the precision of the haptic device or the Nano17 force sensor (US-6-2 ATI standard calibration, which corresponds to 6 lbf maximum in the shear directions). Furthermore, we can visualize force tracking performance as in Fig. 3C. The desired X-and Y-direction velocity commands (received via UDP from the iBCI control computer) are plotted with their corresponding shear force commands as computed via the velocity-shear saturation function (Eqn. 5). In addition, the raw measured shear force is plotted.

#### 2.6.2 Closed-loop iBCI Cursor Control Task

The general iBCI decoding system is composed of two parts taken from machine learning decoding techniques: (1) Open-loop training (or decoder calibration), where a probabilistic model of neural responses is trained on a data set of simultaneously recorded movements and associated neural activity, and (2) Closed-loop control, where the model constructed on the open-loop training data is used in real time to map neural activity patterns to estimated movement trajectories, which can then control computer cursors or robotic manipulators (e.g., [39]–[41]).

All open-loop tasks resembled a standard radial 8 autonomous cursor-to-target-and-back trajectory, with participant T5 being cued to attempt directional hand movements about the wrist joint in concert with the cursor. Specifically, a cursor (45 pixels in diameter) would travel autonomously from the center of the workspace (1920 pixels wide by 1080 pixels tall) to a radial target at one of eight equally spaced locations that were 409 pixels from center. The computer monitor was positioned approximately 75 cm from T5’s eyes. For each trial, the cursor would start at center and begin to move towards a random target after 500 ms. T5 remained idle when the cursor was at center and did not attempt any movement. The cursor’s movement duration was exactly 1.2 seconds to enable T5 enough time to recognize the cursor’s movement and execute the associated attempted movement. All attempted movements were wrist pointing, which we will refer to as *Attempted Hand Joystick*. After the 1.2 second travel duration, the cursor would remain at the target location (where T5 was instructed to hold the attempted movement) for a hold time of 500 ms. After the hold time, the cursor would return to the center of the workspace with the same movement duration of 1.2 seconds, while T5 would make the associated attempted hand movement back to the idle position. There was no haptic feedback given during these open-loop tasks.

For each closed-loop iBCI cursor control session, T5 performed a series of cursor-to-target-and-back blocks that alternated between the *Haptics* condition and the *No Haptics* condition. The cursor-to-target-and-back task does not at any point reset the cursor or end-effector to the center position. Instead, the participant controls a cursor from a center target out to radial targets and back during the entire block. Figure 2 illustrates the system diagram of the iBCI cursor control system integrated with the Phantom Premium haptic device. Each session began with a 3-minute practice open-loop block during which T5 was able to familiarize himself with the *Attempted Hand Joystick* movement strategy. After the practice period, one 4-minute open-loop *No Haptics* block of data was collected to calibrate an initial decoder (Kalman velocity filter [39], [42]). After the initial decoder build, two or three sets of four 5-minute closed loop blocks were performed. Normally, each set comprised two *Haptics* blocks and two *No Haptics* blocks, the order of which was randomly determined prior to each session. A total of seven online sessions were run over the course of six consecutive weeks. Two sessions were run each week with the exception of two weeks where no sessions were run. The alternating (A-B-A-B) paradigm was conserved within all sessions. Due to within-day non-stationarities in neural recordings [43], the decoder was recalibrated after every set, where the previous set’s data from successful trials (2 blocks of *Haptics* and 2 blocks of *No Haptics*) was used as training data for recalibration. Each decoder was built using all available (192) channels.

Task parameters for closed-loop cursor control differed from the open-loop task. Specifically, the cursor was smaller (25 pixels in diameter), the target was smaller (80 pixels in diameter), and the target hold time was longer (700 ms). Target hold time is defined as the continuous duration of time the cursor must remain within the target to register an acquisition of that target. A trial was considered unsuccessful if the cued radial target was not acquired within 10 seconds. These parameters were tuned for moderate to high task difficulty to keep T5 from reaching a performance ceiling such that differences in performance between conditions may be observed. These parameters were found during a pilot study prior to the main study. During closed-loop control, T5 was also urged to keep his head and neck still to mitigate any movement against the tactor as this would result in shear forces not aligned with the commanded shear forces. To aid in this, T5’s head movement was spatially tracked via an OptiTrack V120:Trio camera system which measured the position and orientation of a worn headband outfitted with reflective markers. The head tracking system mapped the coronal plane position of the worn headband to an on-screen green “head cursor”, which visually informed T5 if his head was moving excessively.

Performance within each day was assessed between the *Haptics* and *No Haptics* conditions. Our primary task performance measure was time to target, defined as the time between target onset (when the target appeared) and when the cursor entered the target prior to target acquisition (i.e., time to target did not include the 700 ms hold time necessary for target acquisition; as introduced in [39]). Time to target is only defined for successful trials (approximately 90% of trials in these sessions). We also excluded trials immediately following a failed trial, since the cursor starting position was not the previous target and could be close to the current trial’s target. Other performance metrics computed were path efficiency (a value between 0 and 1, with 1 being a direct straight line) and target dial-in time (time from when the cursor first enters the target to when the target is acquired). All statistical analysis was performed using the Wilcoxon signed-rank test.

## 3 RESULTS

### 3.1 Sensitivity Test Results

Figure 4 summarizes sensitivity to touch for the control group and T5 at probed locations above the upper torso. For the control group, average sensitivity (reported in units of gram-force, or gf) at each location is reported with 95% confidence intervals (Fig. 4B). Figure 4A shows each enumerated probing location, mapping the average sensitivities to their corresponding classification color. The control group has sensitivity classified as *normal touch* (detection of touch less than 0.15 gf) at most probed locations. A few locations, predominantly in the upper back area below the neckline, were classified as *diminished light touch* (detection of force between 0.16 and 0.5 gf). Classifications of either *normal touch* or *diminished light touch* are considered to be in a healthy range. T5 had similar sensitivity to the control group above the neckline (*normal touch*), but, sensitivity immediately degraded below the C4/C5 dermatomes – the location of T5’s spinal cord injury. T5’s sensitivity was classified as diminished protective sensation (detection of force between 0.6-3 gf) in the upper chest area and *loss of protective sensation* (detection of force greater than 4 gf) in the upper back area. Given these results, we identified the back of the neck as the best location to provide haptic feedback.

### 3.2 Results for Perception of Skin-shear Stimulation

#### 3.2.1 Validation of Shear Force Stimuli

The Phantom Premium haptic device is natively optimized for a particular workspace defined by the configuration in which the stylus is perpendicular to the base (e.g., when the stylus is held like a pen). Due to limitations in mounting the device, we used it in the configuration depicted in Fig. 2. Since force production capabilities at the end-effector change as a function of the device’s linkage configuration, we sought to characterize the device’s performance in producing the set of shear force stimuli used during the perception task.

Figure 5 and Table 1 summarize the Phantom Premium’s performance in producing shear force stimuli in 8 radial directions on the back of T5’s neck. We analyzed force sensor measurement data from one session of the perception study. For each trial, we obtained the force sensor measurement at the final time step (the 1 second mark) of the step input of shear force, assuming that this represented the time when the stimulus had reached steady state. Clustering each of these steady-state force vectors into groups by their respective stimulus direction, we computed means and 95% CIs for both force magnitude and direction for each stimulus grouping. Additionally, we computed the mean and 95% CIs of direction error and magnitude error across all trials (i.e., across all stimulus directions).

**Fig. 5:**
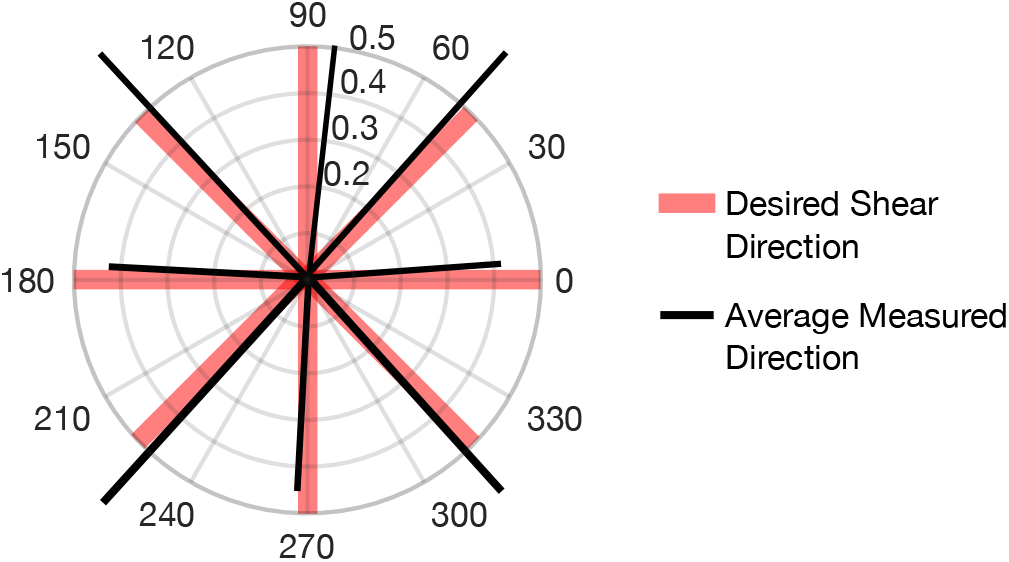
Validation of shear force stimuli. The Polar plot depicts radial directions in degrees where each ring represents a force magnitude in units of Newtons. Values of the average measured directions and magnitudes are provided in Table 1.

**Table 1.**
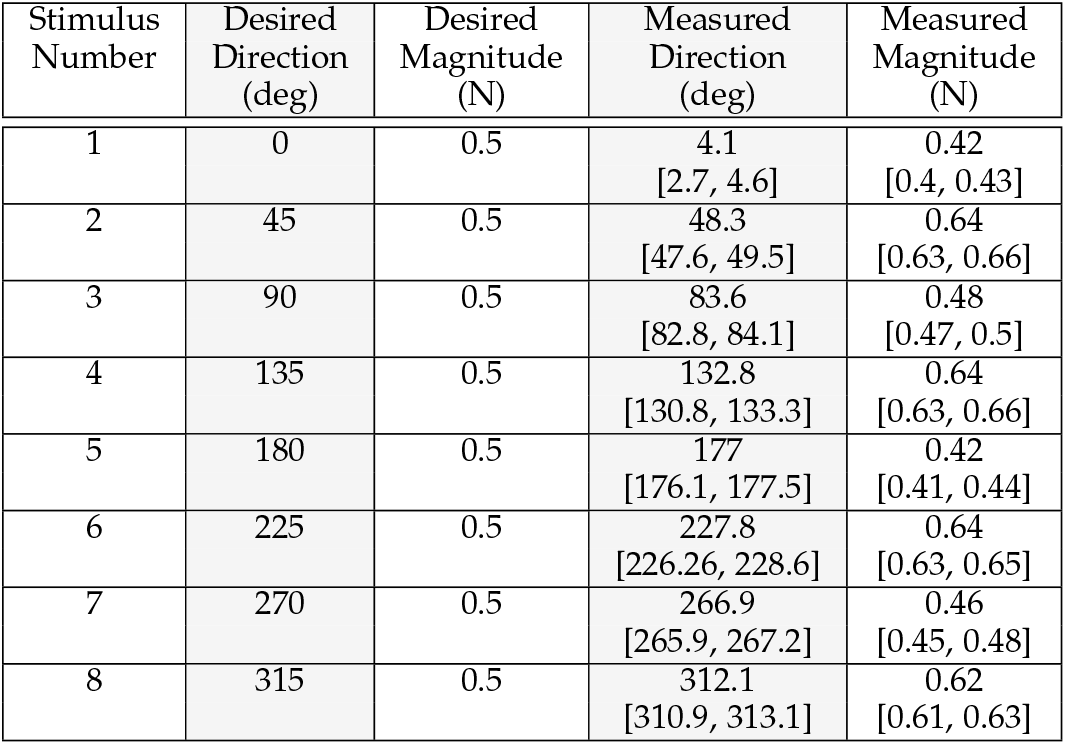
Summary of measured shear force stimuli. Means and 95% CIs (within square brackets) are reported for each stimulus direction.

| Stimulus Number | Desired Direction (deg) | Desired Magnitude (N) | Measured Direction (deg) | Measured Magnitude (N) |
| --- | --- | --- | --- | --- |
| 1 | 0 | 0.5 | 4.1<br>[2.7, 4.6] | 0.42<br>[0.4, 0.43] |
| 2 | 45 | 0.5 | 48.3<br>[47.6, 49.5] | 0.64<br>[0.63, 0.66] |
| 3 | 90 | 0.5 | 83.6<br>[82.8, 84.1] | 0.48<br>[0.47, 0.5] |
| 4 | 135 | 0.5 | 132.8<br>[130.8, 133.3] | 0.64<br>[0.63, 0.66] |
| 5 | 180 | 0.5 | 177<br>[176.1, 177.5] | 0.42<br>[0.41, 0.44] |
| 6 | 225 | 0.5 | 227.8<br>[226.26, 228.6] | 0.64<br>[0.63, 0.65] |
| 7 | 270 | 0.5 | 266.9<br>[265.9, 267.2] | 0.46<br>[0.45, 0.48] |
| 8 | 315 | 0.5 | 312.1<br>[310.9, 313.1] | 0.62<br>[0.61, 0.63] |

Results indicate that the haptic device was fairly accurate in terms of stimulus direction with an overall mean absolute angular error of 4.1°, with 95% CI [3.6°, 4.2°]. Angular accuracy was better in some directions than others, with the largest average angular error of approximately 6.4° occurring for Stimulus 3 (the 90° direction). Figure 5 indicates no structure or systematic offset present in angular errors across each stimulus direction (i.e., there is no constant rotational error between desired and measured stimulus directions). Although angular errors exist, the errors are smaller than the angle between each stimulus direction (which was 45°).

In terms of force magnitude, the haptic device had an overall mean absolute magnitude error of 0.1 N, 95% CI [0.09 N, 0.11 N] across all stimulus directions when the desired force magnitude was 0.5 N. Force magnitude in the diagonal directions (approximately 0.6 N) were greater than the force magnitude in the cardinal directions (approximately 0.45 N) on average. We believe this was a byproduct of the haptic device’s kinematics at the configuration used. Additionally, the errors in direction could also be due to anisotropy of the skin stiffness in different directions [44], [45]. Nonetheless, the average shear force magnitude in each direction was equally perceivable to T5; he mentioned that all stimuli had the same force.

#### 3.2.2 Perception of Shear Direction

Figure 6A summarizes T5’s perception of shear force direction on the back of the neck in the form of a confusion matrix. Out of 488 total trials (61 repetitions for each stimulus), T5 predicted 320 trials correctly, for a classification accuracy of 65.6%. Erroneous classifications were predominantly made between adjacent shear directions, as illustrated by the color banding about the diagonal axis in the matrix. T5 was most accurate in classifying the 90° and 270° stimuli, with accuracy of approximately 80%. T5 was most inaccurate in classifying shear stimuli in the diagonal directions of the lower hemisphere (the 225° and 315° directions), where he tended to misclassify them as their adjacent horizontal directions (i.e., 180° and 0°, respectively).

**Fig. 6:**
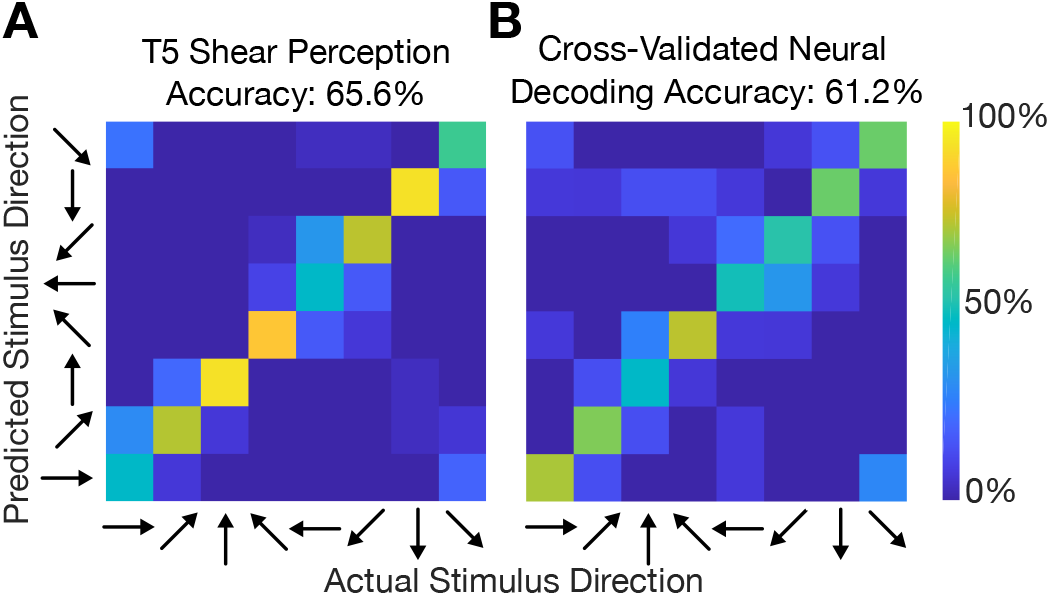
Cognitive perception of sensory stimulus versus decoding sensory stimulus from neural firing rates. (A) Confusion matrix of participant T5’s shear direction perception. (B) A Gaussian Naïve Bayes classifier was used to classify each trial’s stimulus using firing rates within the 500 to 900 ms window after stimulus onset. Only channels which significantly responded to shear stimulation were used (21 channels). For each matrix, the entry (i,j) in the matrix is colored according to the percentage of trials where stimulus j was decoded/predicted (out of all trials where stimulus i was cued).

### 3.3 Tactile Responses in Motor Cortex

We found some electrode channels that were visibly modulated by the tactile shear stimulus, as seen in the peristimulus time histograms (PSTHs) in Figure 7. Threshold crossing spike firing rates (mean ± 95% CIs) are shown for four example electrode channels across all 8 shear stimulus directions. Some electrode channels, e.g., Channel 153, have clear responses after stimulus onset, as indicated by the increase in firing rate from baseline within the window during which the shear stimulus was provided.

**Fig. 7:**
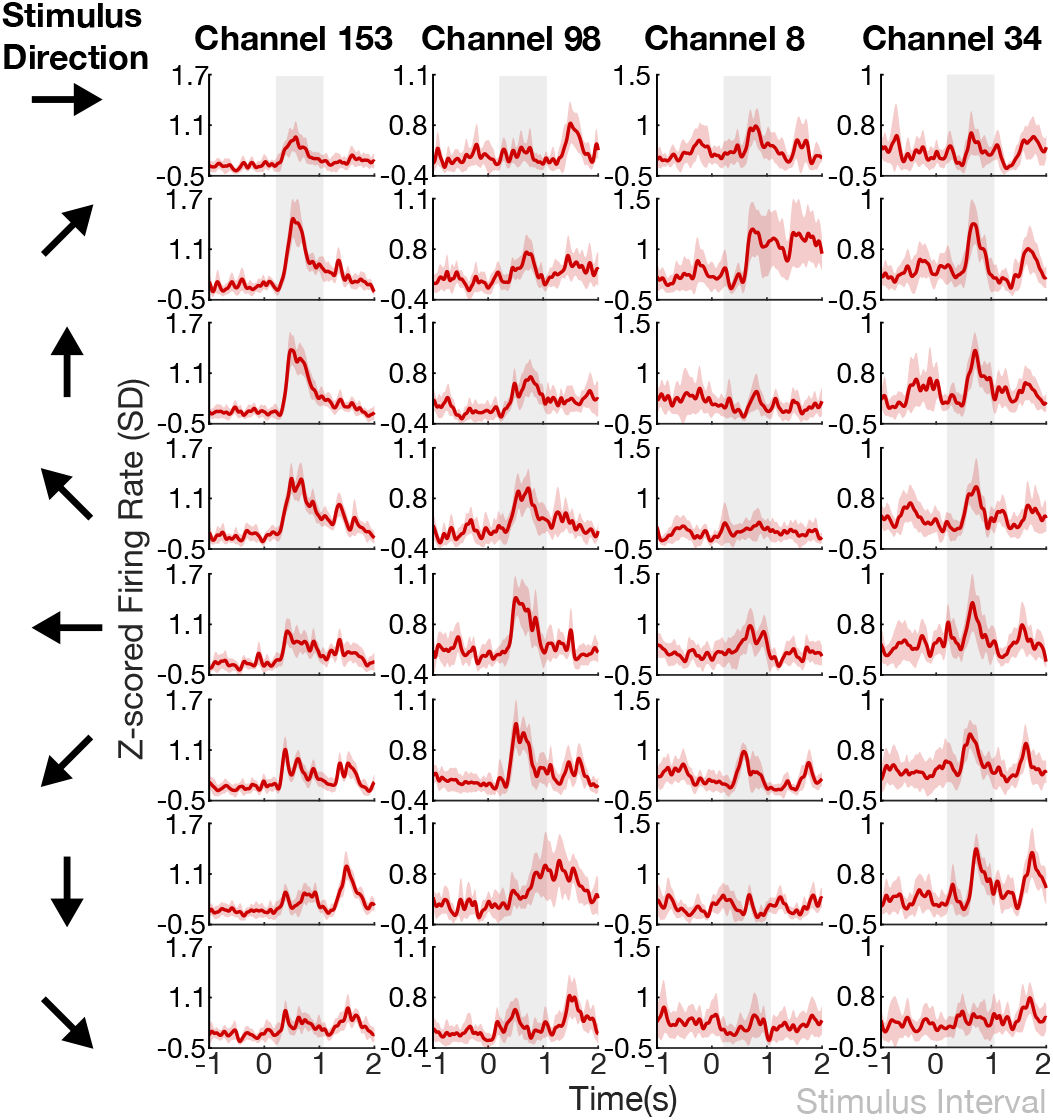
Peristimulus time histograms, shown as red traces for four example electrode channels (columns) across skin-shear direction (rows). Firing rates (mean ± 95% CIs) and the window during which shear stimuli were provided (gray) as a step input are shown.

Electrodes with significant tuning to shear stimulation are depicted in Figure 8. Approximately 21 out of 95 of functioning electrode channels were found to be significantly modulated by skin-shear stimulation on the back of the neck. Significant modulation is defined as a significant difference in firing rates between the idle state activity and activity measured during the stimulation period as computed via 1-way ANOVA. Furthermore, we found a range of tuning strengths across all significantly tuned electrodes, reported in FVAF scores. A high FVAF score indicates that the particular electrode responds differently to each stimulus direction, whereas lower FVAF scores mean that the electrode responds to each stimulus direction in a similar manner. The distribution of significantly tuned electrodes indicate no somatotopic or orderly organization of stimulus preferences across the hand knob area of the motor cortex, but rather an even scattering across arrays.

**Fig. 8:**
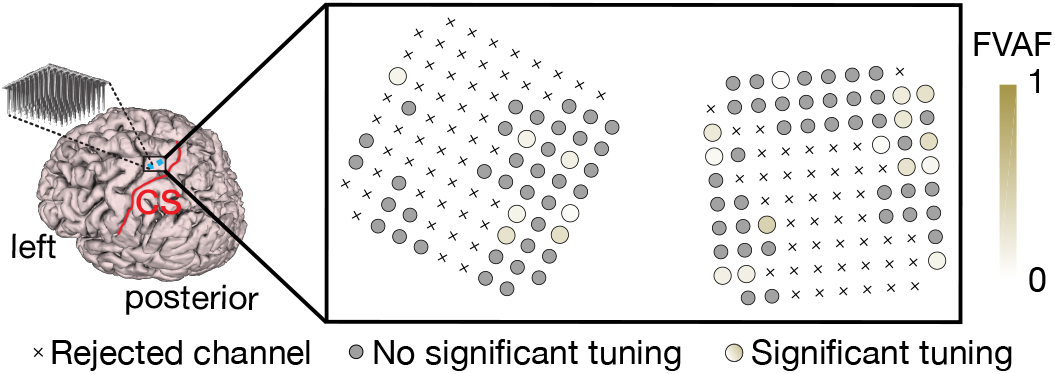
Shear-related tuning strength across electrode arrays. The strength of each electrode’s tuning to shear stimulus is indicated with a shaded gold color (darker indicates more tuning). Tuning strength was quantified by computing the fraction of total firing rate variance accounted for (FVAF) by changes in firing rate due to the stimulus directions. Crosses represent “non-functioning” electrodes. Small gray circles indicate channels with no significant tuning to shear stimulation. Larger colored circles indicate significantly tuned channels.

Figure 9 depicts tuning curves computed for 11 example electrode channels significantly tuned to shear stimulation. The tuning curves exhibit different shapes, but curved shapes were observed more often than linear ones. To assess how this tuning compares to the known cosine directional tuning of motor cortical cells [46], we fit the curves with a cosine-tuning model and report the distribution of R^2^ values for the 21 significantly tuned electrode channels. We found that the R^2^ values for over half of the tuned channels were greater than 0.5, indicating that these channels generally have preferential tuning to a particular stimulus direction with tapering firing rates as the stimulus direction rotates away.

**Fig. 9:**
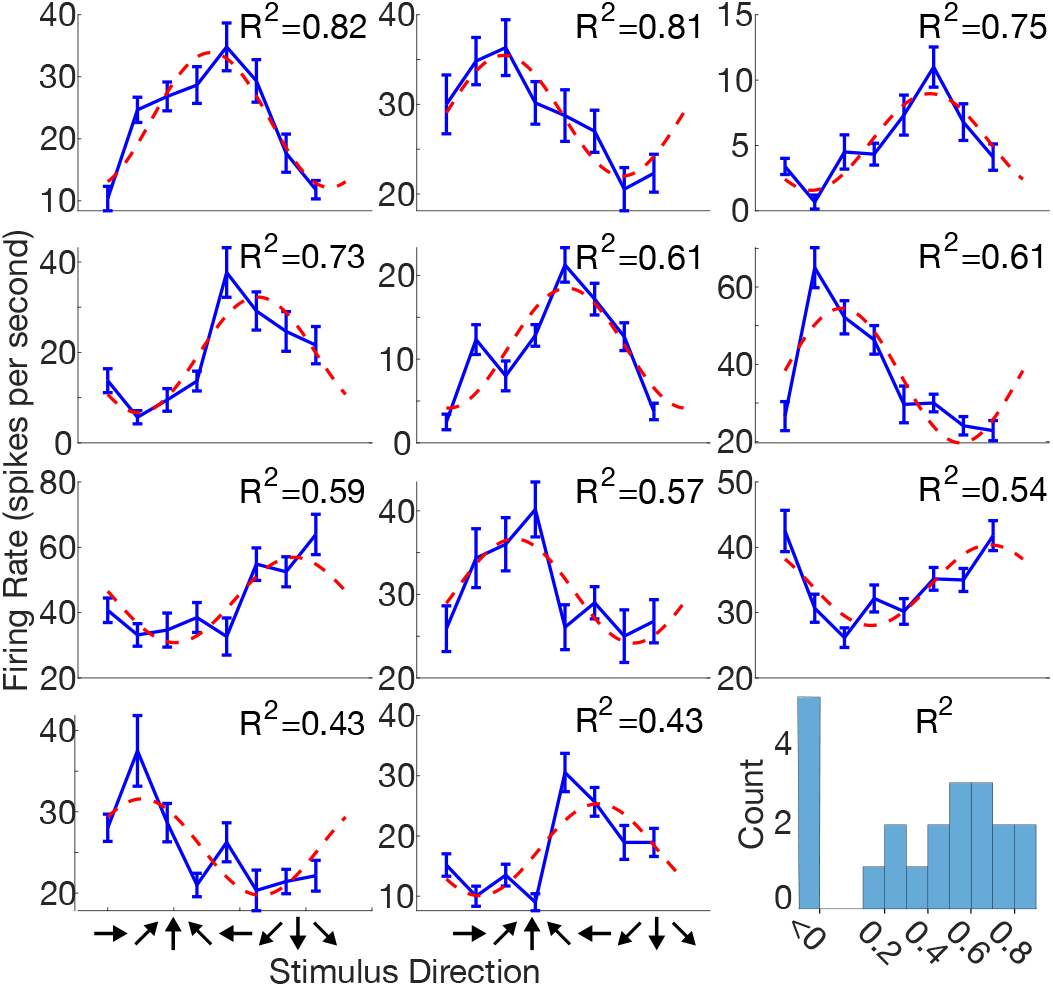
Tuning curves for shear stimulation. Example tuning curves show the range of shapes observed across significantly tuned electrode channels. Each panel corresponds to a single exemplary electrode channel. Blue curves indicated mean threshold crossing firing rates within a 500 to 900 ms window after stimulus onset with linear interpolation; error bars show standard error of mean. Red curves are fits to a cosine-tuning model with corresponding adjusted R^2^ values shown in the upper-right corner of each panel. Bottom right: adjusted R^2^ histogram for all 21 significantly tuned electrode channels. Values less than 0 indicate that a linear fit is better than the cosine-tuning model.

Observing the depth of tuning to shear stimulation across significantly tuned electrode channels, we applied an offline leave-one-out cross validated Naïve Bayes decoder to classify the shear direction on a given trial based on threshold crossing firing rates within the 500 to 900 ms window after stimulus onset. The decoder achieved a classification accuracy of 61.2% (Fig. 6B) using only the 21 significantly tuned electrodes. This classification accuracy based on neural firing rates is similar to T5’s verbal classification accuracy (65.6%) of stimulus direction during the perception task discussed in Section 3.2.2.

### 3.4 iBCI Cursor Task Performance With and Without Haptic Feedback

We examined the Radial 8 cursor control task data to determine what effect the haptic feedback had on cursor task performance. As shown in Figure 10A, the task was performed as a sequence of blocks (alternating red and gray clusters of points) during which participant T5 either was or was not given skin-shear haptic feedback acting as artificial proprioception. Within-session time to target performance is detailed in Table 2. Time to target performance was only significantly better for three out of the seven total sessions (*p* ≤ 0.5 for two sessions, *p* ≤ 0.01 for one session). The overall across-session median time to target was 4.12 seconds (3.45 s ± 1.99 s mean ± s.d.) for the *Haptics* condition and 4.24 seconds (3.56 s ± 1.99 s mean ± s.d.) for the *No Haptics* condition. Time to target performance during the *Haptics* condition was slightly but significantly (*p* ≤ 0.05) better than the *No Haptics* condition across all sessions. Figure 10B depicts the median within-session time to target, path efficiency, and dial-in time as a function of session number. The median time to target and median dial-in time for the *Haptics* condition were lower than the *No Haptics* condition in each session except the first. Likewise, median path efficiency was greater for the *Haptics* condition in each session except the first. Path efficiency and dial-in time performance for the *Haptics* condition was slightly but significantly (*p* ≤ 0.01) better than the *No Haptics* condition in session 5.

**Fig. 10:**
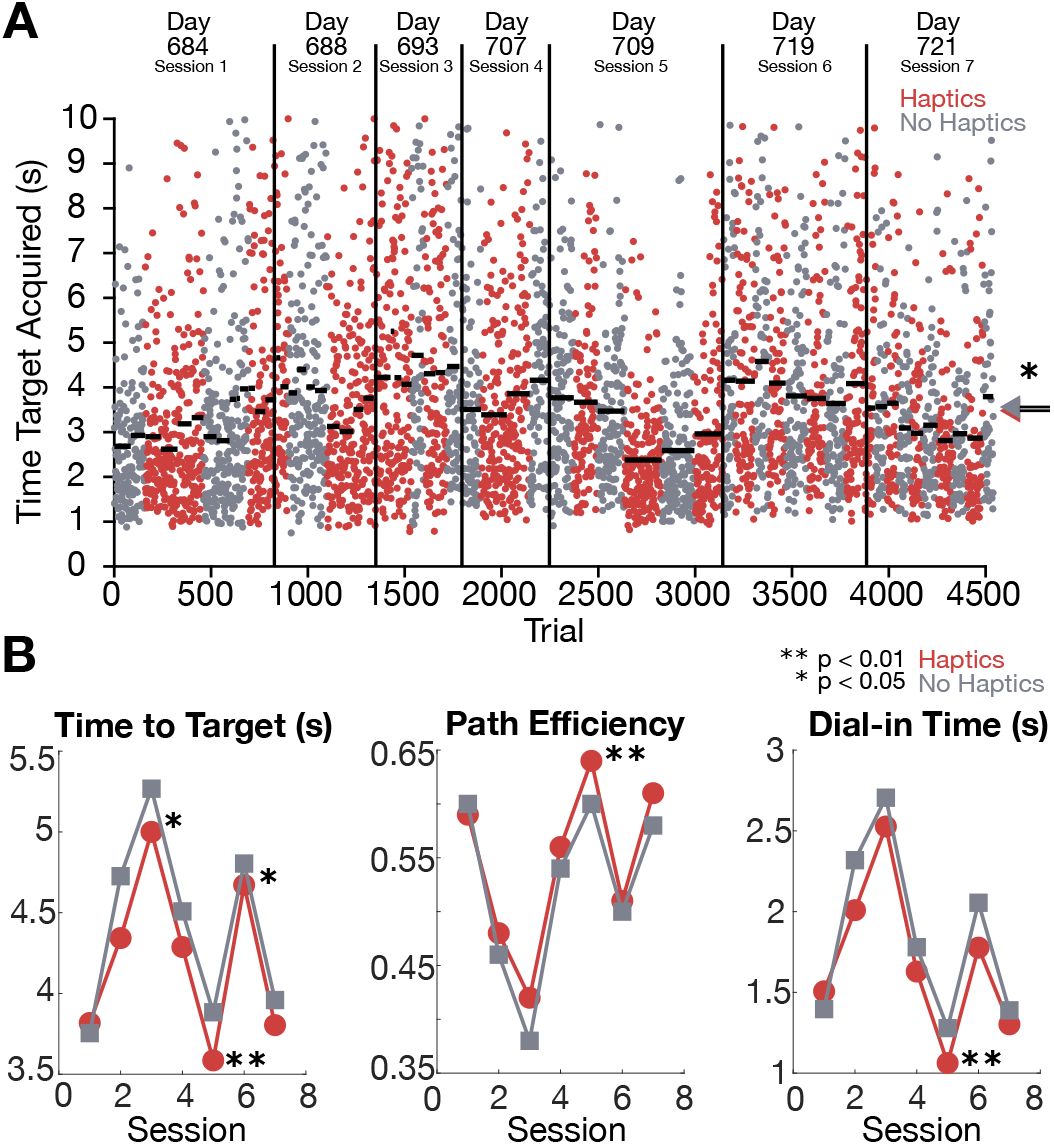
iBCI cursor control performance with and without haptic feedback. (A) Timeline of iBCI comparison blocks across 7 sessions (day numbers are days after implantation). Red clusters indicate *Haptics* blocks where the participant received skin-shear haptic feedback, and gray clusters indicate *No Haptics* blocks where haptic feedback was disabled. Each dot shows one trial’s time to target. Horizontal bars show the median time to target of each block. Arrows on the right show the median across all trials of each condition. (B) Median time to target, path efficiency, and dial-in time within each session is plotted for each condition. Asterisks indicate statistical significance level: * p≤ 0.05, ** p≤ 0.01.

**Table 2.** Summary of iBCI cursor control performance. Median time to target is reported with means and standard deviations inside parentheses. P-values computed using the Wilcoxen signed-rank test are reported with asterisks indicating level of significance: * *p ≤* 0.05, ** *p ≤* 0.01.

| Session Day | Haptics Condition Time to Target (s) | No Haptics Condition Time to Target (s) | p-value |
| --- | --- | --- | --- |
| 1 | 3.82 (3.13 ± 1.87) | 3.76 (3.08 ± 1.80) | 0.73 |
| 2 | 4.34 (3.67 ± 2.05) | 4.72 (4.05 ± 2.24) | 0.04* |
| 3 | 5.00 (4.32 ± 2.35) | 5.27 (4.59 ± 2.52) | 0.31 |
| 4 | 4.29 (3.62 ± 1.96) | 4.51 (3.83 ± 1.99) | 0.20 |
| 5 | 3.59 (2.91 ± 1.65) | 3.88 (3.20 ± 1.70) | 0.002** |
| 6 | 4.67 (3.99 ± 2.03) | 4.80 (4.13 ± 2.12) | 0.49 |
| 7 | 3.80 (3.12 ± 1.82) | 3.96 (3.28 ± 1.96) | 0.03* |
| **Overall** | **4.12 (3.45 ± 1.99)** | **4.24 (3.56 ± 1.99)** | **0.012\*** |

## 4 DISCUSSION

In this study, we demonstrated the existence of a population of cortical units in the hand knob area of the premotor cortex that are tuned to tactile sensory inputs derived from shear forces applied to the back of the neck of a person. Additionally, we assessed our participant’s perception of shear force in 8 unique directions on the back of the neck and compared his verbal classification accuracy (65.6%) to that of a linear classifier decoding a subset of neural activity (21 electrode channels) resulting from shear force stimuli. Finally, we designed, implemented, and demonstrated online iBCI cursor control with an integrated desktop haptic feedback device driven by iBCI-decoded velocity commands. Time-to-target performance was slightly but significantly better with haptic feedback than without (*p* ≤ 0.05).

The motor cortical responses to shear force stimuli were distributed across the microelectrode arrays with no apparent somatotopy. The tuning characteristics were well modeled by cosine-shaped functions, where neural activity was modulated enough that a linear classifier was able to decode the stimulus direction from threshold crossing firing rates. The correct classification rate was approximately 61.2% over 8 possible stimuli, which is on par with previously reported classification rates with human ECoG participants between 52.4 and 66.7% [22] over 3 possible stimuli. We used a simple linear decoder (Naïve Bayes) to classify average firing rates within a time window of interest and since the neural activity was time varying, as seen in the PSTHs in Figure 9, more sophisticated nonlinear decoders including recurrent neural networks [47], multiscale dynamical models [48], and sequential autoencoders [49] may have the potential to decode with higher accuracies.

To gain insight as to whether we should have expected the neural population (21 significantly tuned channels) decoding accuracy (61.2%) to have exceeded the participant’s verbal perception accuracy (65.6%), we can look to studies concerned with which components of neural activity evoked by a sensory stimulus is meaningful for perception. This field often compares behavioral psychometrics to “neurometrics” – functional relationships that express the sensitivity of neuronal responses to a sensory stimulus or motor behavior [50], [51]. Early pioneering studies in the middle temporal (MT) cortical areas, involved with visual motion analysis, suggest that the stimulus discrimination and detection capacity of single sensory neurons can be close to or even surpass the perceptual ability of the animal [50], [52], [53]. However, more recent studies from various groups suggest that firing-rate-based neurometric performance (both in single neuron analysis and population-level analysis) does not exceed perceptual performance [54], [55], and instead is more similar to one another where the perceptual performance acts as a type of upper bound that the neurometric performance tends towards [56], [57]. One main caveat is that these previous studies measured neural activity in areas of the brain that do not overlap with the hand knob area of premotor cortex that we measure from in this study. However, the studies suggesting that neurometric performance tends towards perceptual performance help provide insight as to why our neural decoding accuracy may have fallen short of the participant’s perceptual accuracy.

Past literature has documented somatosensory responses in motor cortical units, although these studies have mostly reported responses to passive movements of limbs [58]–[63]. Many of these studies conceptualized these results within the framework of a “reflex” that is receiving local muscle spindle information about the perturbed joint and activating muscles to generate corrective movements. Only recently, an ECoG study with human participants has reported sensory responses, in the finger/hand area of the motor cortex, to light passive brushing of finger digits [22]. However, this study also noted similarity in responses to proprioceptive finger bending, ultimately believing it unlikely that a sensory-specific subpopulation was recorded.

A key differentiating element of our study from previous literature is the location on the body where the haptic stimulation was applied. Haptic stimulation was provided on the back of the neck, whereas neural activity was measured from the so-called ‘hand knob’ area of the motor cortex (pre-central gyrus). One could assert that haptic shear stimulation at the back of the neck should not elicit a reflex response from the hand and/or arm and thus should not result in sensory responses in the motor cortex. However, if we consider the recent findings of whole-body representation in this same small patch (‘hand knob’ area) of the motor cortex (pre-central gyrus) [23] – including the head and neck – then neck-related reflexes due to shear stimulation may be possible and potentially lead to the sensory responses recorded. A post-assessment of session videos for the perception tasks indicated little to no movement of the head in response to the shear stimulation because the head was firmly resting (lightly pressed) against the headrest at all times, greatly reducing the potential for the measured responses to be related to head movement.

Results from the closed-loop iBCI control experiment indicate that haptic feedback does not interfere with or degrade iBCI control based on decoding attempted hand movements. In fact, haptic feedback led to a significant though small improvement in time to target performance. The small improvement in performance during the haptic feedback condition could either be due to (1) the participant’s ability to cognitively perceive and use the additional feedback to better accomplish the task, while the relatively small size of neural modulation to haptic feedback did not significantly influence the decoder, or (2) the possibility that the haptic feedback positively influenced the decoder. If the former is true, then the results of this study add to a growing body of work indicating that velocity decoding is robust to other processes reflected in motor cortical activity, such as visual feedback [64], object interaction-related activity [65], or other concurrent motor tasks [66]. Recent reports also indicate that, when attempting to move multiple body parts concurrently, the “dominant” body part suppresses neural activity related to the the others [23], where the ‘dominant’ body part is the one most highly represented amongst the sampled population of neurons. The ‘dominant’ body part is the arm/hand in our case, so arm/hand related neural activity could have potentially suppressed the sensory responses to the concurrent haptic feedback, resulting in little influence to the iBCI decoder during control. The relatively small size of neural modulation, along a dimension coding for haptic feedback, also provides insight as to why the haptic feedback may not have influenced the decoder greatly.

Now, let us consider the case that haptic feedback could have potentially influenced the decoder positively. One theory is that the haptic feedback could have boosted the signal-to-noise ratio if the tuning curves for the motor cortical sensory responses are aligned with the tuning curves for directional attempted hand movements. Alternatively, the haptic feedback could have contributed to the participant engaging in attempted movements with greater intensity, as prompted by the forces, which theoretically would have been decoded as velocity commands of greater magnitude. This may have resulted in the small improvement in time to target performance for the *Haptics* condition, although a follow-up study would be necessary to thoroughly investigate this hypothesis. It is likely that the haptic feedback had little influence on the iBCI decoder, and that the participant was able to utilize the feedback as an additional information channel, which led to the marginal increase in performance. Interestingly, the participant did not find the haptic feedback device distracting, in fact, he mentioned that it provided him with a sense of “feeling the cursor move”.

At the time of this study, there was very limited literature regarding touch sensitivity using the SWME across the body in people at locations other than the hands and feet [67], [68]. Bell-Krotoski et al. [69] used the same SWME kit to assess sensitivity of a healthy population in a few locations along the arms, legs, and face. Their results are consistent with ours in similar probing locations, including locations 6 and 14 of the upper chest and locations 3 and 7 of the face. Although the SWME kit is a convenient and easily deployable clinical test, it is highly subject to human error. Care must be taken by the study operator in administering the exam, specifically monitoring the angle and speed of monofilament application. In a pilot study, we found that higher-velocity probes are more likely to elicit a response due to the excitation of higher-frequency modes upon contact between the filament and skin. It is unclear as to which mechanoreceptors, the biological sensors that detect tactile stimuli, are the target of the SWME since the frequency content of probing forces cannot be easily measured. To mitigate the chance of introducing bias in touch sensitivity evaluation, higher-fidelity testing may be achieved with haptic devices which are often capable of delivering finely controlled forces with high accuracy and precision (e.g., 3D Systems Phantom Premium, Force Dimension omega.3). Nonetheless, this SWME kit was used to assess simple touch sensitivity to help define a target location on the body, upon which we could provide a higher-fidelity haptic stimulation using a desktop Phantom Premium haptic device.

Using the haptic device, we demonstrated that participant T5 can perceive the direction of a 0.5 N shear force stimulus provided on the back of the neck in one of eight radial directions with an accuracy of 65.6%. Interestingly, T5’s perception of shear direction was worst in the lower diagonal directions where he would commonly perceive them as the closest horizontal direction. Barring these lower diagonal directions, T5’s classification accuracy amongst the other 6 directions was roughly 80%. At first, we considered that T5’s perception of the lower diagonal directions may have been diminished by a nearby surgical scar just below the area of stimulation. However, upon further investigation, we realized that if the scar was interfering with perception, then it would have also interfered with perception of the 270° stimulus as well. This was not the case. Another reason may be attributed to the mechanical properties of the skin in different planar directions, as previous studies have reported nonlinear stiffness properties of the glabrous skin under tangential shearing [45].

A group has previously conducted a relevant but limited skin-shear perception study on the fingertip, where participants were able to distinguish between four shear directions separated by 90° [70]. Another group reported that participants were incapable of distinguishing between ‘slip’ stimuli – sliding across a surface of the skin – within 20° of one another on the fingertip. Although these studies were performed on the fingertip, the results give us an idea of perception of skin-shear direction on the back of the neck, giving us a potential lower and upper bound on perceptual limitations of shear direction (i.e., between 20° and 90°). It is important to reiterate that the aforementioned skin-shear and skin-slip studies have been applied to the fingertip, which is known to have a high density of mechanoreceptors [71]; the density and distribution of types of mechanoreceptors in the hairy skin on the back of the neck is unclear. To our knowledge, this is the first study of its kind to assess perception of skin-shear on the back of the neck. We did not perform the shear perception study on an able-bodied control group, as our goal was to ultimately incorporate shear force feedback into an iBCI control task for participant T5. Therefore, we only assessed T5’s perception of shear.

In addition to assessing T5’s perception of shear force, we characterized the performance of the Phantom Premium device in producing a set of shear forces while in a non-optimal configuration. The overall angular error in each direction was well below the resolution of stimulus directions. The force magnitude varied at an average of 0.1 N from the desired 0.5 N. Drawing again from skin-shear literature on the fingertip, it is believed that approximately 0.28 N of shear force is necessary to convey direction of shear with high accuracy, assuming fingertip skin stiffness of 1.4 N/mm and displacement of 0.2 mm [70]. The stiffness of the hairy skin on the back of the neck is presumably less than that of the glabrous skin on the fingertip, so the threshold for shear detection is likely well below 0.28 N on the back of the neck. Therefore, we can assume that force magnitudes of 0.5 ± 0.1 N are well above the 0.28 N conservative threshold for direction detection, and the 0.1 N average error lies below the detectable threshold.

There are a few limitations to this study that should temper over-generalization of these interpretations. First, the study was conducted with a single participant. Additionally, the participant was quite familiar with 2-dimensional iBCI cursor control, having approximately 2 years of experience at the time of this study. We attempted to increase the difficulty of the task by tuning parameters motivated by Fitts’ Law such as the target diameter, target distance, cursor diameter, velocity gain, and target hold time, but, it is possible that the participant reached a performance ceiling where we would not be able to see a greater difference in task performance between the tested conditions. And finally, random haptic perturbations would allow better understanding of the influence of motor cortical responses to sensory stimulation on decoding during online cursor control. Specifically, investigating whether iBCI control performance degrades if random haptic perturbations occur which are not aligned to cursor kinematics or the target directions. This would help elucidate whether the haptic feedback positively influenced neural activity during iBCI control or if it merely made the participant more attentive and alert resulting in higher firing rates as seen in previous literature when recalibrating decoders using closed-loop data [37], [39], [40], [72].

## 5 CONCLUSION

The results of this study suggest that peripheral haptic feedback may be a viable method of communicating task-relevant information during BCI control that does not impose additional loads on the predominantly used visual sensory channel. Because peripheral haptic stimulation can elicit natural sensations by leveraging intact sensory pathways at locations of the body with intact sensitivity, it may be easier to learn how to decipher the information encoded in the stimulus, as opposed to learning how to decipher the sensations elicited via intracortical microstimulation. Future studies are needed to quantitatively compare these approaches [18]. Also, other decoded parameters could be used to drive haptic feedback to communicate other task-relevant information, such as position, which could potentially be used in the absence of vision to convey spatial information of a cursor’s location. Additionally, peripheral haptic feedback could be used to communicate interactions with objects when controlling prostheses.

## Acknowledgements

We, most importantly, thank BrainGate clinical trial participant T5 who tirelessly dedicated his time and energy to participate in these studies. We also thank Beverly Davis, Erika Siauciunas, and Nancy Lam for administrative support. DRD was supported in part by the National Science Foundation Graduate Research Fellowship Program and in part by the Stanford University Bio-X Graduate Fellowship Program. PR was supported in part by NIH-NIDCD R01-DC014034. LRH was supported in part by NIH-NIDCD R01DC014034, NIH-NINDS UH2NS095548; Office of Research and Development, Rehab. R&D Service, Department of Veterans Affairs (B6453R, N2864C), The Executive Committee on Research (ECOR) of Massachusetts General Hospital, and the MGH Deane Institute for Integrated Research on Atrial Fibrillation and Stroke. KVS and JMH were supported in part by NIDCD R01-DC014034, NIDCD U01-DC017844, NINDS UH2-NS095548, NINDS UO1-NS098968, Larry and Pamela Garlick, Samuel and Betsy Reeves, the Wu Tsai Neurosciences Institute, and the Bio-X Institute at Stanford University. KVS was supported in part by Simons Foundation Collaboration on the Global Brain 543045 and Howard Hughes Medical Institute (Investigator) at Stanford University.

